# Light environment drives the shallow to mesophotic coral community transition

**DOI:** 10.1101/622191

**Authors:** Raz Tamir, Gal Eyal, Netanel Kramer, Jack H. Laverick, Yossi Loya

**Author notes:** Corresponding author: Raz Tamir.

## Abstract

Light quality is a crucial physical factor driving coral distribution along depth gradients. Currently, a 30 m depth limit, based on SCUBA regulations, separates shallow and deep mesophotic coral ecosystems (MCEs). This definition, however, fails to explicitly accommodate environmental variation. Here, we posit a novel definition for a regional or reef-to-reef outlook of MCEs based on the light vs. coral community-structure relationship. A combination of physical and ecological methods enabled us to clarify the ambiguity in relation to that issue. To characterize coral community structure with respect to the light environment, we conducted wide-scale spatial studies at five sites along shallow and MCEs of the Gulf of Eilat/Aqaba (0-100 m depth). Surveys were conducted by Tech-diving and drop-cameras, in addition to one year of light spectral measurements. We quantify two distinct coral assemblages: shallow (<40 m), and MCEs (40-100 m), exhibiting markedly different relationships with light. The depth ranges and morphology of 47 coral genera, was better explained by light than depth, mainly, due to the Photosynthetically Active Radiation (PAR) and Ultra Violet Radiation (1% at 76 m and 36 m, respectively). Branching coral species were found mainly at shallower depths i.e., down to 36 m. Among the abundant upper mesophotic specialist-corals, *Leptoseris glabra*, *Euphyllia paradivisa* and *Alveopora* spp., were found strictly between 36-76 m depth. The only lower mesophotic-specialist, *Leptoseris fragilis*, was found deeper than 80 m. We suggest that shallow coral genera are light-limited below a level of 1.25% surface PAR and that the optimal PAR for mesophotic communities is at 7.5%. This study contributes to moving MCEs ecology from a descriptive-phase into identifying key ecological and physiological processes structuring MCE coral communities. Moreover, it may serve as a model enabling the description of a coral zonation world-wide on the basis of light quality data.

## Introduction

Coral reefs constitute spectacular and diverse marine ecosystems. As reef-building corals maintain a mutualistic symbiosis with photosynthetic dinoflagellates (Trench 1993), light intensity and spectral quality play an important role in successful coral colonization (Frade et al. 2008). The light spectrum affects the initial stages of coral settlement, with planulae exhibiting species-specific responses (Mundy and Babcock 1998). Additionally, the light regime is a key factor for various stony corals at advanced life stages, in determining their survival and growth (Mass et al. 2007, Lesser et al. 2009). Consequently, light can have a substantial effect on coral (Vermeij and Bak 2002, Hennige et al. 2010).

Light conditions, in both the ultraviolet (UVR: 290–400 nm) and photosynthetically active radiation (PAR: 400–700 nm) spectra, undergo change with increasing depth. These changes are largely a function of the optical properties of the water itself (Kirk 2011). The absolute amount of downwelling irradiance (*Ed*) decreases with depth. The spectral composition also becomes increasingly dominated by the ultraviolet/blue part of the spectrum (Kirk 2011). This environmental gradient is an important factor in controlling the productivity, physiology, and ecology of corals (Gattuso et al. 2006, Frade et al. 2008, Cooper et al. 2010, Kahng et al. 2010, Ben-Zvi et al. 2015) with different species necessarily occupying different depth ranges or niches (Kahng and Kelley 2007, Bridge et al. 2012, Eyal et al. 2015). Even slight and temporary changes in water transparency can exert a crucial effect on mesophotic reefs operating near the limits of photosynthesis (Kahng et al. 2010). Therefore, varying light conditions may affect how coral communities are structured in space (Kahng et al. 2010, Bauman et al. 2013).

The euphotic zone ends at the depth where 1% of surface PAR remains (Z_1%_) (Kirk 2011). Given the relationship of stony corals with their photosynthetic endosymbionts, Z_1%_ is expected to have a notable influence on the depth distribution of various species, despite some exceptional corals capable of flourishing deeper than Z_1%_ (Fricke and Meischner 1985; Schlichter et al. 1994; Lesser et al. 2009; Pochon et al. 2015). The attenuation coefficient of downwelling PAR (K_d(PAR)_) describes the optical nature of water, and relates directly to the euphotic zone according to the Lamberte Beer law (I_z_ = I_0_ e^-Kdz^) (Kirk 2011). Light penetration into a water body linearly correlates with optical water quality (Gattuso et al. 2006, Kirk 2011). Therefore, even when light at the surface is equal for two locations, its quality (i.e. intensity and spectrum) may differ at the same depths at those locations due to changes in K_d(PAR)_. Kahng *et al*. (2010) demonstrated the potential of K_d(PAR)_ to explain variation in the depth limits of zooxanthellate corals among locations worldwide. Moreover, the influence of light attenuation on the distribution of other light-dependent marine organisms has also been documented at local scales, albeit on much smaller scales (Manuel et al. 2013).

Though PAR enhances coral growth at certain depths, light can also have negative impacts. Under high light levels there is the potential for damage to the holobiont (bleaching) (Dunne and Brown 1996). The biologically damaging effects of UVR are well known, and include the direct effects of UVB (Holm-Hansen et al. 1993) and possible effects mediated through reactive oxygen species (ROS) (Mallick and Mohn 2000, Tchernov, Kvitt et al. 2011). Dunne & Brown (1996) tested the penetration of solar UVB radiation in shallow tropical waters and calculated DNA-damage. Their function returned a threshold for DNA damage at 1% surface UVB. Therefore, the 1% limit of both PAR and UV is expected to have a crucial effect on coral distribution patterns.

Mesophotic Coral Ecosystems (MCEs) flourish under limited light, and at greater depths than shallow reefs (Lesser et al. 2009). Despite the relatively large area occupied by MCEs (Eyal et al. 2015, Lesser et al. 2018), and a recent surge in research attention (Menza et al. 2007, Laverick et al. 2018), biogeographical and ecological data are sparse. Strong gradients exist in downwelling solar irradiance on mesophotic reefs (Lesser et al. 2009), and a broader understanding of the abiotic factors (Lesser et al. 2018) may help to explain and predict patterns in coral-reef communities along spatial and vertical scales.

Despite their close proximity to well-studied shallow reefs, and their inferred importance (Rocha et al. 2018), MCEs have remained poorly studied due to the technical limitations of diving (Pyle 2019). Basic data on the taxonomic composition, depth ranges, habitat preferences, abundance and distribution of MCE taxa are scarce. Moreover, the processes that structure these communities are virtually unknown (Hinderstein et al. 2010). One of the main outstanding issues is that of how species-specific responses to differing light conditions influence the transition from shallow to mesophotic communities. Edmunds *et al*. (2018) noted that “given the importance of light in the ecology of coral reefs, it may be timely to reconsider the value of high-resolution in situ sampling for this parameter for time-series analyses of coral reefs”. Moreover, Lesser *et al*. (2018) concluded that “to improve the current definition of MCEs, which may result in regional or reef-to-reef definitions, we need more studies that include community characterization throughout the entire depth range of 30–150 m, that are combined with studies on the optics of the water column”.

In this study, we sought to address this knowledge gap. We provide ecological data pertaining to stony coral reefs across depth and space in the Gulf of Eilat/Aqaba (GoE/A), together with light quality data. The present work emphasizes the crucial importance of light regimes to the structure of coral communities in both vertical and horizontal space, while offering a novel model for coral reef zonation that combines physical and ecological methods.

## Materials and Methods

### Data collection

Spatial benthic surveys were conducted over 10 km of reef at five sites along shallow reefs to MCEs (0-100 m depth). Belt-transects, 50 m in length, were recorded in bins between 5-100 m (5, 10, 20, 30, 40, 50, 60, 70-80, 90-100 m) parallel to the shore at each site. 2,320 photo-plots (70×50 cm) from all transects were analysed for live coral cover and bathymetric substrate type (e.g. rock, gravel and dead corals) and non-available settlement area (sand). We used the multiple points method in the ‘CoralNet’ web interface (Available at: http://coralnet.ucsd.edu/) to estimate percentage cover in the photo-plots. Thirty photo-plots were randomly selected from each 50 m transect to record coral genus abundance, for later community structure analysis.

Light and water temperatures were recorded monthly from August 2014 to July 2015, using a profiling reflectance radiometer (PRR800, Biospherical Instruments, USA), which measures 19 channels (at 300–900 nm) and the integrated PAR. The instrument was deployed at midday (11:00–13:00), using the free-fall technique (Waters et al. 1990) to maintain a vertical position and avoid shading and reflectance from the boat. The measurement data were analysed using the program PROFILER (Biospherical Instruments, USA). An average depth for 1% irradiance of surface UV and PAR, and 0.1% PAR, and PAR attenuation coefficient (K_d_PAR) were calculated (from 8-19 different GPS locations), as described by Kirk (2011). 14 years of daily Chlorophyll-*a* concentration data were provided by the national monitoring program (NMP) in Eilat from 01/01/2004 - 31/12/2017 (Available at: http://www.meteotech.co.il/EilatYam_data/ey_observatory_pier_download_data.asp).

### Statistical analyses

The corals identified from 40 independent transects were treated as samples. Sample matrices were log (1+x) transformed to Bray-Curtis dissimilarity matrices. Analysis of Similarity (ANOSIM; 999 permutations) was used to test for differences in a multivariate coral community matrix using pairwise dissimilarities. SIMPER analysis was used to determine which coral genera contributed to the differences observed between depths (Clarke 1993). Analyses were performed at the genus level using PRIMER-7 software. To visualise dissimilarities, a non-metric multidimensional scaling (nMDS) plot was created in R (Team 2013) by Oksanen *et al*. (2013). Additionally, Species richness and Hill number of Shannon diversity index (Sahnnon 1948) were calculated per depth.

We identified co-occurring assemblages of coral genera using a recently published approach derived from a mesophotic system in the Caribbean (Laverick et al. 2017). We performed a Principal Coordinate Analysis (PCoA) on Hellinger transformed data to enable a method based in Euclidean space. Data for each species in a photo-plot were standardised by total abundance across all photo-plots (Legendre and Gallagher 2001). The Euclidean distances between taxa in community space can be considered as measures of dissimilarity. We used K-means clustering (Hartigan et al. 1979) to determine the optimal number of communities to fit to the PCoA. We trialled clustering solutions fitting 2-10 assemblages and selected the number of assemblages that maximised the Calinski criterion (Caliński and Harabasz 1974). The returned assemblages were tested statistically with a multi-response permutation procedure (MRRP) running 9,999 iterations (Biondini et al. 1988). We performed a Dufrene-Legendre indicator species analysis (D-F, 10,000 iterations) (Dufrêne and Legendre 1997) in the R package (Roberts 2016) on the photo-plot data. Our D-F analysis returned how strongly each photo-plot resembled any assemblage identified by the PCoA with K-means clustering. Maximum indicator values occur when all Scleractinia observed in a photo-plot are from a single assemblage. The depth ranges of taxa forming each assemblage were compared in a t-test to determine whether this depth range structured the reef community. Depth ranges were limited to the 10^th^-90^th^ percentile of a taxon to remove the influence of extreme observations.

To assess the influence of light on community structure, DF indicator values for each assemblage at each photo-plot were plotted against %PAR and %UV, pooled across sites. Non-linear least squares models were fitted to the data. A measure of model parsimony (Akaike information criterion; AIC) was used to determine the best model for explaining the ecological data based on depth, %UV or %PAR, and monthly light data. To contextualise the light data, a time series decomposition was performed using the package series (Trapletti et al. 2015) on 14 years of Chlorophyll-*a* data. The seasonal effects were extracted to characterise the expectation for the annual spring phytoplankton bloom in the GoE/A.

## Results

Patterns in depth distributions for different genera and species were found to be correlated with light metrics, i.e. UVR and PAR (1% at 36 m and 76 m, respectively – Fig. 1). Similarly to previous works (Laverick et al. 2017) we found on more than one occasion that communities identified as either shallow or mesophotic could be vertically separated by only 10 m of water, demonstrating remarkable changes in community structure as a result of minor changes in depth. Overall, there were significant differences in community structure between depths (ANOSIM, R^2^ = 0.76, p < 0.05), but not among all five sites (ANOSIM, R^2^ = 0.061, p > 0.05).

**Fig. 1.**
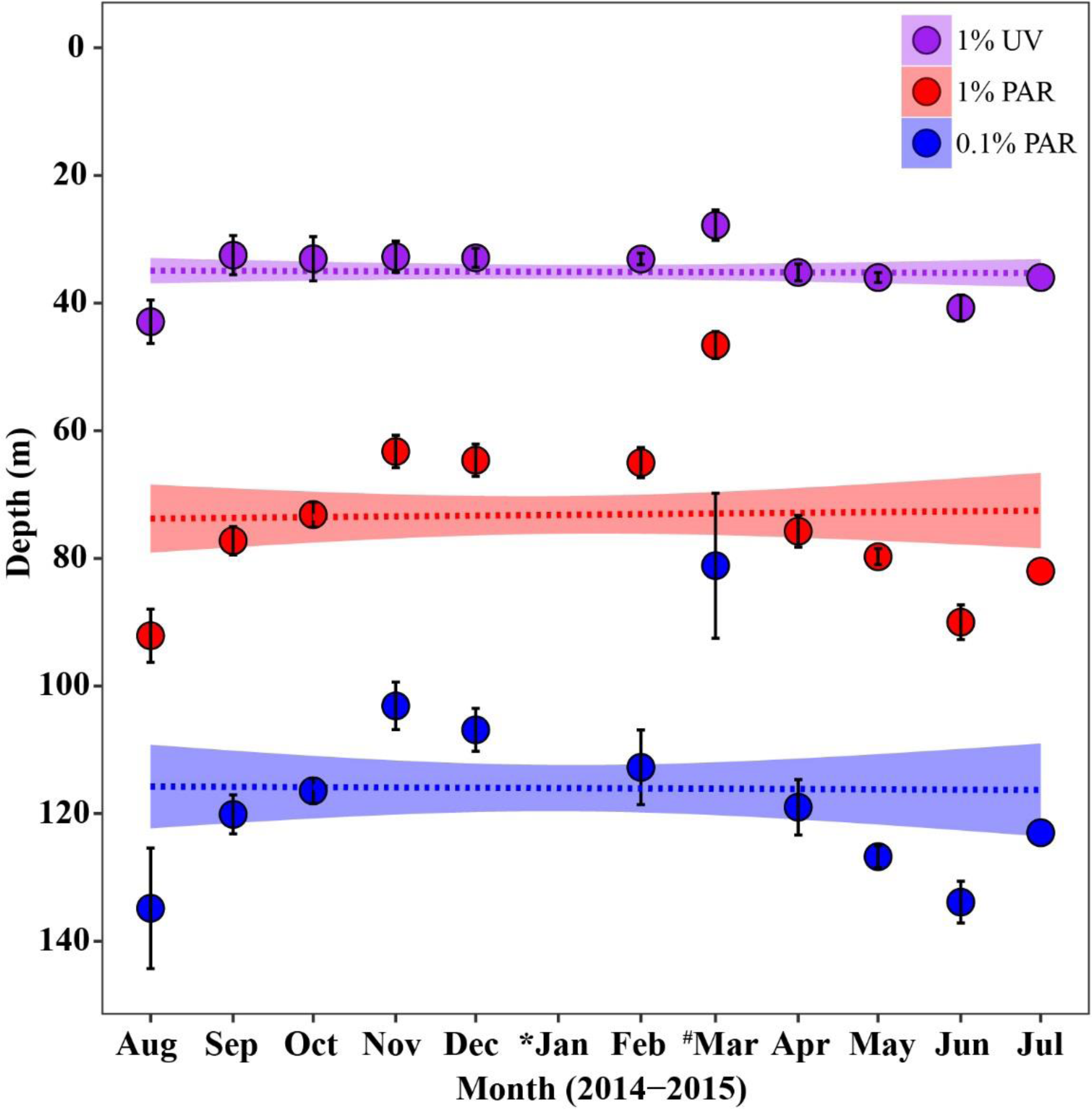
Annual light penetration through depth, averaged over eight different sites (bottom depth deeper than 200 m) offshore Eilat. Dots represent the mean depth of 1% irradiance of surface UV light (purple – 36 m), 1% PAR (red – 76 m) and 0.1% PAR (blue - 119 m) for each month with CI bars; horizontal lines denote the annual average value, with CI area (transparency). *(January) indicates no data, and ^#^(March) indicates the range of measurements from March 2015 (affected by an algal bloom event). July was measured once.

Patterns in coral zonation are correlated with coral morphology, i.e. branching species dominate shallow depths (< 40 m) (Fig. 2a-d) and foliaceous species dominate 40-80 m depth (Fig. 2i-l), with a massive coral threshold at 70m depth (Fig. 2e-h). The spatial ordination of coral community structure revealed that the largest changes occurred at depths of 40 and 70 m (Fig. 2 & Table 1S). Branching coral species are shallow-specialists, found mainly above 40 m (Figs. 2a-d & 1S). *Acropora* spp. and *Stylophora pistillata,* the most common branching corals at the GoE/A (Fig. 1S), were found in limited abundance deeper than this line, with a reduction of 67% (*Acropora* spp.) and 96% (*S. pistillata*) in the number of colonies between 40 to 50 m. None of those genera were found deeper than 60 m. The maximum depth observed for the third most common branching coral, *Seriatopora hystrix,* was 40 m.

**Fig. 2.**
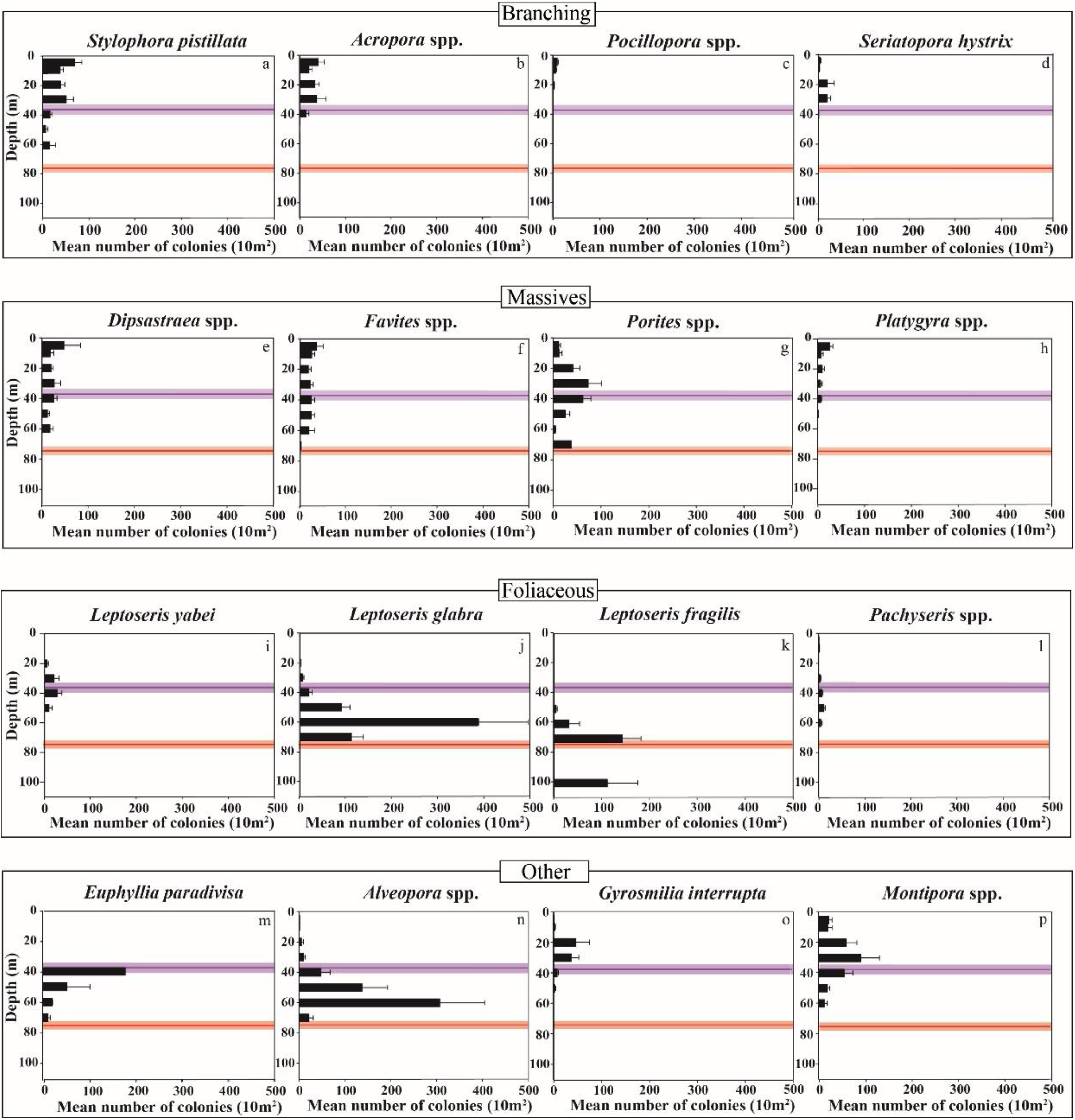
The mean (±SE) number of different species/genera of scleractinian coral colonies along a depth gradient of five survey sites. The purple line represents the annual average of 1% UVR (36 m depth) with 95% confidence interval (purple area). The red line represents the annual average of 1% PAR (76 m depth) with 95% confidence interval (red area).

Overall, we found a clear change in community structure at 40 m depth. Abundant corals, such as *Porites* spp., *Montipora* spp., *Favites* spp. and *Dipsastraea* spp. were found to flourish down to the maximum 1% PAR limit. Among the abundant upper mesophotic specialist-corals, *Leptoseris glabra*, *Euphyllia paradivisa* and *Alveopora* spp., were found mostly between 36-76 m depth (Fig. 2 m, n, o). *L. glabra* was highly abundant at 60 m (1160 colonies, 386.6 ± 109 mean *(± SE)* per site). Only a few colonies (27 colonies, 6.75 ± 2.5 mean *(± SE)* per site) of this species were found at 30 m depth, the minimum depth of this species, and no colonies were observed below 70 m depth.

The strongest indication of a light-limited depth range can be seen in the distribution of *E. paradivisa,* which was observed strictly between the 1% UV and 1% PAR depths. *E. paradivisa* was restricted to 36-76 m. In addition, clear changes in coral abundance along the depth gradient were found (Fig. 1aS). The transition from ‘shallow’ 0-40 m, which is dominated by branching and encrusting genera to ‘deep – upper mesophotic’ 50-70 m, dominated by foliaceous genera, mostly *Leptoseris* spp. and ‘other’ genera such as *Alveopora* spp. (Fig. 1S). The ‘lower mesophotic’ >70 m was dominated by *Leptoseris* spp. and azooxanthellate corals such as *Rhizopsammia* and *Dendrophyllia* spp. (Fig. 1aS). Supporting these patterns, our SIMPER analysis returned a clear signal in community structure across the different depth zones (SIMPER and ANOSIM pairwise tests, global R^2^ = 0.833, P < 0.001, stress = 0.07) ‘shallow’ (≤40 m), ‘upper-mesophotic’ (50-70 m) and ‘lower-mesophotic’ (≥ 70 m) reefs (Table 1S). Further variability is noted between each 10 m depth interval (i.e. 10-20, 30-40, 40-50 m etc.), implying further gradual change (Table 2S).

Overall, sharp changes in stony coral species diversity from the shallow (20-40 m) to the upper mesophotic reef (Fig. 2S), in tandem with changes in light, may indicate changes in physical properties of the water body. Corals found between 20-40 m were thriving reef communities, displaying the greatest species diversity and abundances (Figs. 1S, 2S).

Our analysis further links variability with depth in the light environment to coral assemblages in the GoE/A. Two assemblages of coral genera returned an optimal fit to our data, following PCoA with clustering and a test of the difference between both assemblages (MRRP, P = 0.004, Fig. 3S). The mean depth ranges of each assemblage were not significantly different, with the mean range for clusters 1 and 2 as 30.5 m and 35 m respectively (*t* = −1.04, P = 0.3). Cluster 1 (red) comprises mainly ‘shallow’ genera, with the average cluster 1 photo-plot capturing 8.8 genera (S.D. = 4.7). Cluster 2 (blue) is more varied, but mainly contains ‘deep’ genera, with the average photo-plot capturing 3.3 genera (S.D. = 2.8). Plotting photo-plots spatially, coloured according to the assemblage they most closely match (Fig. 3), further supports one of the assemblages as shallow and the other as mesophotic.

**Fig. 3.**
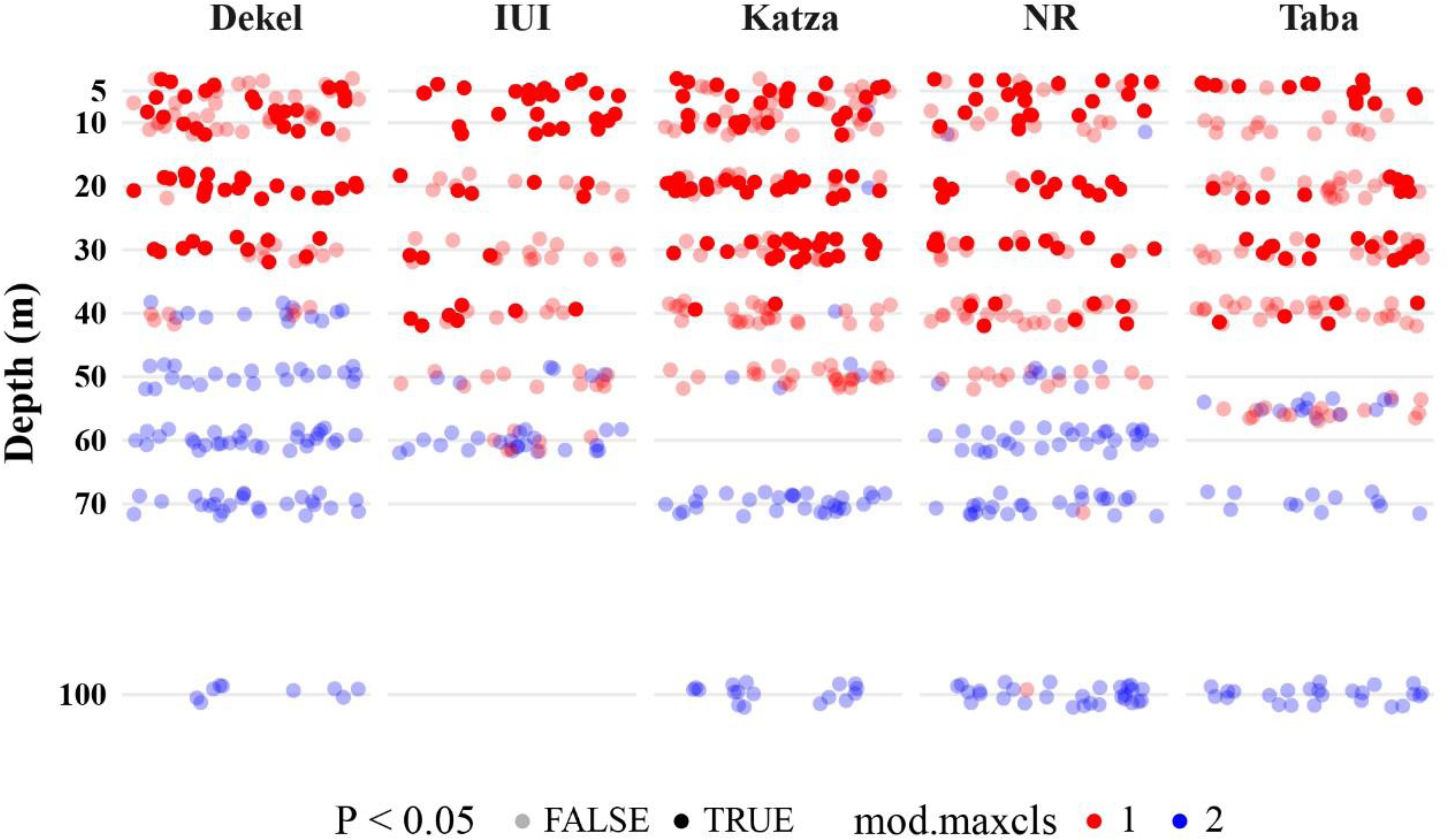
Photo-plots are plotted as points in relation to their depth and site. Points are coloured based on the coral assemblage the plot most closely matches, revealing spatial patterns in assemblages. How closely an observed quadrat aligns with the coral assemblage identified by PCoA analysis was determined by the Defrene-Legendre indicator analysis. Photo-plots are opaque if the P-value returned by indicator analysis is < 0.05; otherwise the point is transparent.

After plotting the D-F indicator values for shallow genera (Cluster 1) we opted to fit a Michaelis-Menten equation (equation 1) (Fig. 4):

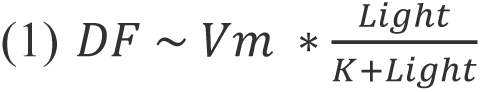

**Fig. 4.**
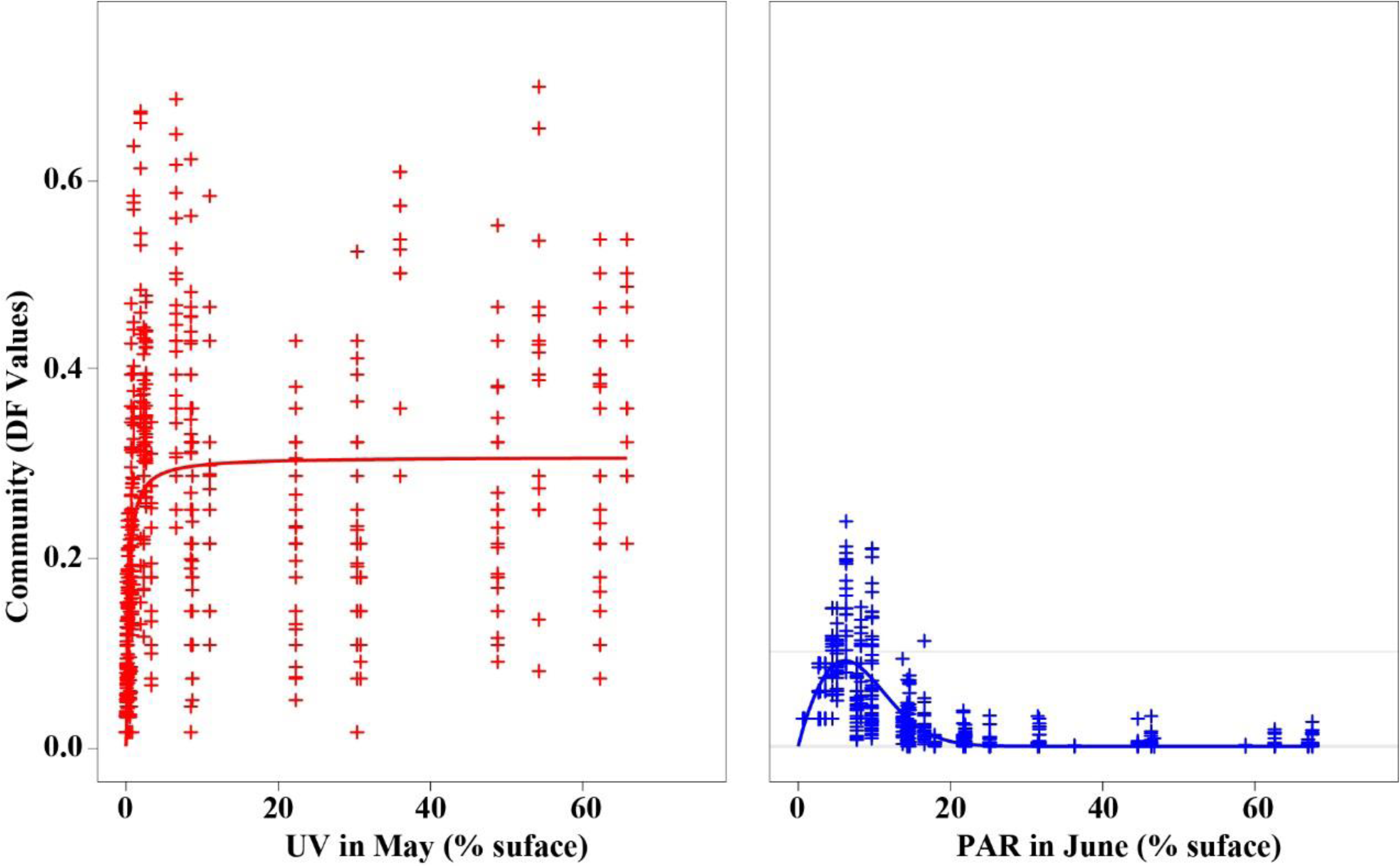
The DF indicator values for cluster 1 (red, shallow) and cluster 2 (blue, mesophotic) are plotted for each photo-plot. The fitted lines show non-linear least squares models after model selection, as described in the results.

May returned the lowest AIC of any month when fitting %UV to data, and March when fitting %PAR (Table 3S). We therefore use these months when evaluating models of DF indicator values against depth, %UV and %PAR. %UV was a better predictor of Cluster 1 D-F values than %PAR, although both light metrics were superior to depth (AIC = −1099, −1034, −666, residual standard error = 0.12, 0.12, 0.15, respectively). Light metrics were able to explain more of the variation in community structure across our data than depth for shallow genera. Vmax (The asymptotic DF value) was estimated as 0.3 ± 0.01 (*t* = 43.39, P < 0.001) for %UV, and 0.3 ± 0.01 (*t* = 38.35, P < 0.001) for %PAR. K (Light required to reach halfway to asymptote) was estimated as 0.29% ± 0.04 (*t* = 7.43, P < 0.001) for %UV, and 1.25% ± 0.15 (*t* = 8.16, P < 0.001).

For mesophotic genera (Cluster 2) we opted to fit a Weibull model (equation 2) (Fig. 9S):

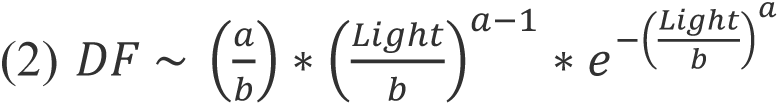

August returned the lowest AIC of any month when fitting %UV to data, and June when fitting %PAR for cluster 2 (Table 3S). %PAR was a better predictor of Cluster 2 D-F values than %UV, returning a lower AIC and presenting a relationship with a mechanistic explanation. Both light metrics returned a lower AIC than depth (AIC = −3249, −2845, −2267, residual standard error = 0.03, 0.03, 0.05, respectively). For %PAR, *b* (shape parameter) was estimated as 9.14 ± 0.16 (*t* = 57.32, P < 0.001), while for %UV *b* = 20.00 ± 1.23 (*t* = 16.26, P < 0.001). *a* (scalar term) was estimated as 1.89 ± 0.05 (*t* = 40.34, P < 0.001) for %PAR, and 0.98 ± 0.01 (*t* = 79.64, P < 0.001) for %UV. The median PAR value returned by the model is 7.5%, or 13.8% UVR, indicating the preferred light environment of the observed mesophotic community.

The light environment of the GoE/A is modulated by an annual phytoplankton bloom. Time series decomposition of Chlorophyll-*a* data reveals that the bloom peaks typically in March (Fig. 5). Note that the seasonality displayed (Fig. 5) is only plotted for three years to improve legibility, but is estimated from all 14 years of monitoring data. The height of the algal bloom in March increases the Chlorophyll-*a* content of the water by more than three-fold in comparison to June through August, and increases the attenuation of light with depth. Key features in this seasonal cycle coincide with the months independently identified as the best light readings to explain coral assemblage distributions.

**Fig. 5.**
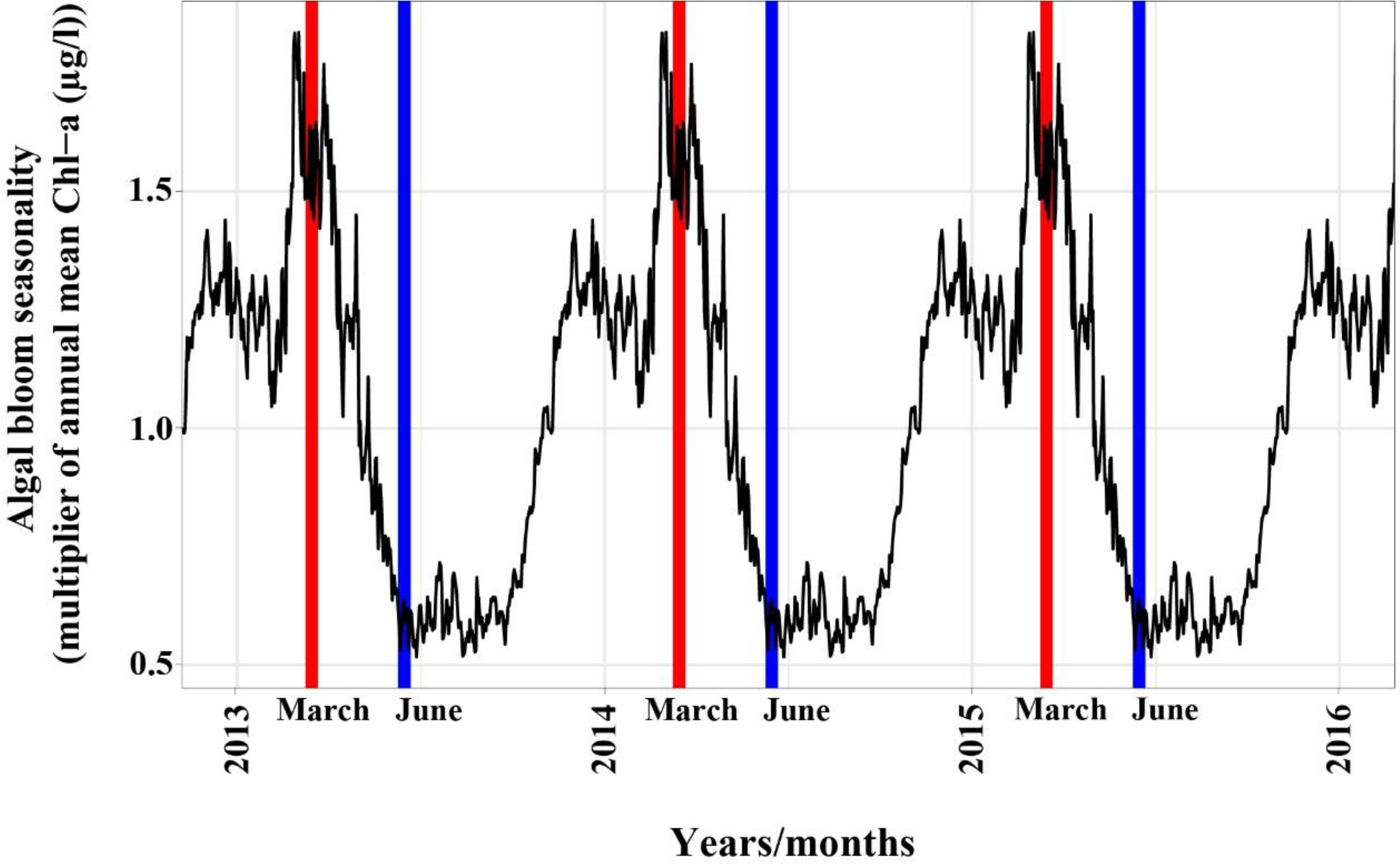
The seasonal variation in ocean Chl-a concentration, calculated from 14 years of monitoring data (black). A value of 1 on the Y-axis is the annual average. The plotted values are the effect of a given month in the year; a month with a value of 0.5 expects a Chl-a concentration of half the annual average. The month of PAR values that best explain shallow (red) and mesophotic (blue) genera distributions coincides with the peak and end of seasonal algal blooms in the northern GoE/A. The values displayed across years are identical, and are repeated to illustrate the cyclical nature of the estimated seasonality of the algal bloom.

### Light attenuation coefficient (Kd_(PAR)_) effect

The varied water characteristics across the GoE/A result in different light attenuation coefficients (Kd_(PAR)_) across sites (Fig. 6a). In parallel to changes in the light attenuation coefficient (Kd_(PAR)_), there is a dissimilarity in coral abundance at the different sites along the GoE/A (Fig. 6b). Overall, the limits of genera and species depth distribution are correlated with Kd_(PAR)_ values. *S. pistillata* is a depth-generalist coral with a depth range of 5-60 m at all the sites, except Dekel Beach, the most light-attenuated site (Kd_(PAR)_ 0.076-0.078 annual average and summer average 0.068-0.07). At this site, *S. pistillata* reached a maximum depth of 40 m. *Montipora* spp. flourished at all depths down to 50-60 m at sites with lower Kd_(PAR)_ values, whereas at Dekel Beach, this species was found at a maximum depth of 40 m and with only five colonies (in 10 m^2^), constituting a reduction of 83% from 30-40 m. We found the same pattern for *Favites* spp., with a maximum depth of 40 m at Dekel Beach and 60 m at the other sites. The mesophotic species *E. paradivisa* and *Alveopora* spp., were the dominant corals of the upper mesophotic zone (40-70 m) except for one site – Katza. *E. paradivisa* at this site dominates between 40-60 m, while at the clearer water site, it flourishes at 60 m and mostly at 70 m. *Leptoseris fragilis* was the most abundant zooxanthellate coral below 70 m depth (Fig. 1S), with no colonies being found shallower than 60 m.

**Fig. 6.**
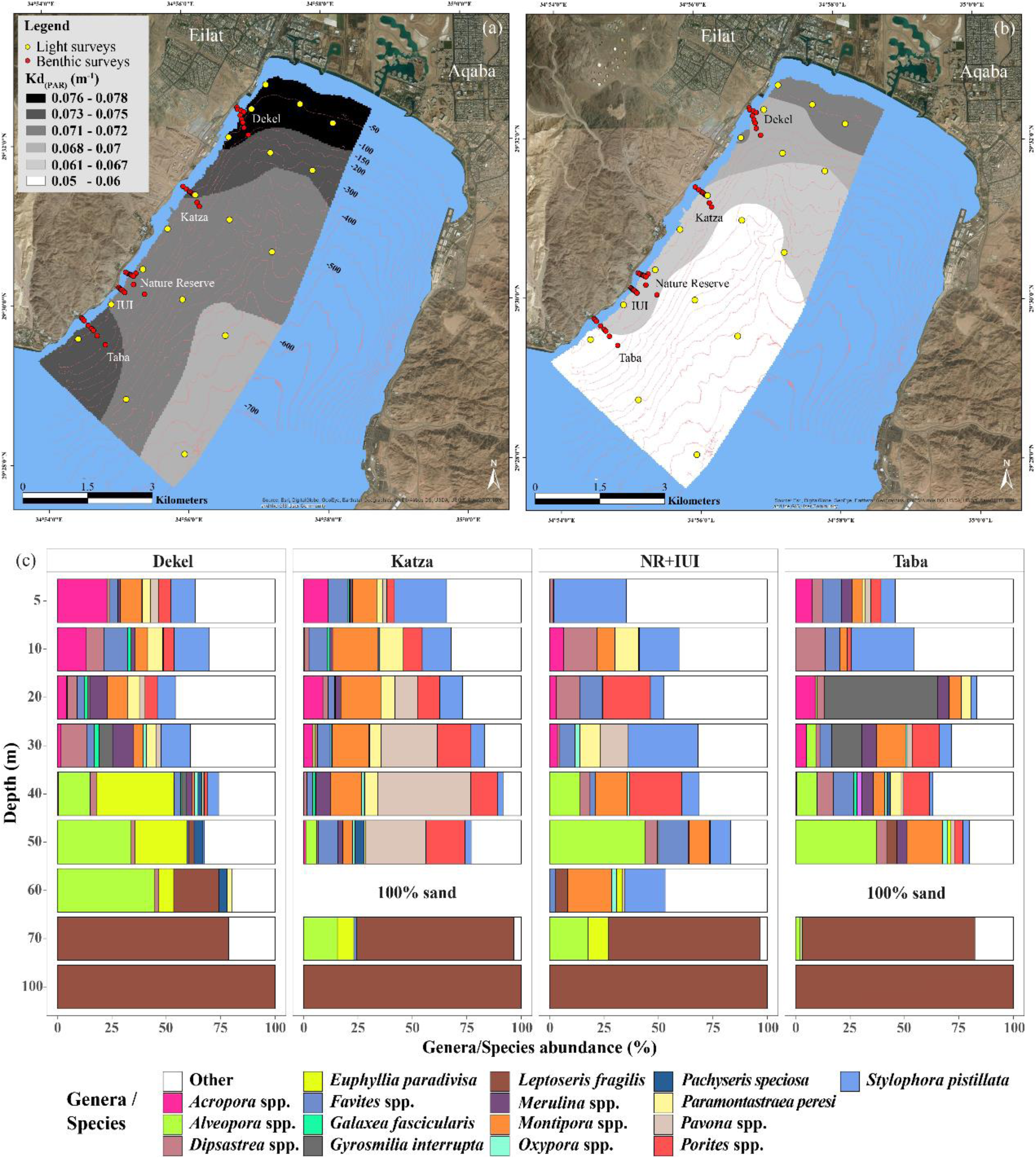
The change in coral community structure as a function of light attenuation among the study sites (a) Annual 2014-2015 average Kd(PAR) values range between 0.076-0.078 (black) and 0.068-0.07 (bright grey) and (b) summer (June-August) average range between 0.068-0.07 (dark grey) and 0.05-0.06 (white) at 30 m depth. Shapefile conducted between the Israeli territorial waters and the 30 m bathymetric depth contour. Yellow dots represent the spectral measurements location. Red points represent transects of all benthic survey site locations. Pink lines represent the bathymetric contours in 50 m intervals. (a-b) Maps were created using ArcGIS Version. 10.2.1 (Esri Inc.) platform. (c) Cumulative percentage of the 17 zooxanthellate corals contributed (threshold - cum. < 50%) to the dissimilarity among the various sites along a depth gradient at the five surveyed sites. IUI + NR sites represent a combination of the two sites. As a result of 100 % sand cover at the Taba and Katza sites at 60 m, no data available.

## Discussion

The survival and growth of coral depends on variable environmental conditions such as light, seawater temperature, nutrient concentration, and salinity among others (Schlichter et al. 1994, Kleypas et al. 1999, Goodbody-Gringley et al. 2015). Still, even though light quality is a crucial physical factor in coral reef ecosystems, little attention has been given to light as a driving force of coral distributions (Edmunds et al. 2018, Lesser et al. 2018). Our results illustrate the importance of understanding light regimes when explaining the distribution and structure of zooxanthellate coral communities, across both depths and spatial gradients. Currently, the distinction between shallow and mesophotic reefs arbitrarily follows SCUBA limitations (Lesser et al. 2018, Eyal et al. 2019, Pyle et al. 2019), or ecological patterns for the lower limit (Semmler et al. 2016, Laverick et al. 2017). We should question, however, whether these approaches can adequately assess between-site variation in rates of community transition with depth (Laverick et al. 2018). We suggest that the light conditions at a given site can be used to explain changes in community structure, when recognised.

### Modern day measurements are consistent with historical studies

Previous studies have documented the photo-adaptive mechanisms of corals, from morphological plasticity to *symbiodinium* density (Hoegh-Guldberg et al. 2007) and pigmentation concentration (Ben-Zvi et al. 2015, Muir et al. 2015), as well as light-enhanced calcification (Goreau 1959, Chalker and Taylor 1975). These mechanisms allow many coral species to maintain their metabolic functions over broad light ranges. Despite such adaptability, coral reefs in general, do have minimum light requirements (Kirk 2011). Limiting-light depths will vary across locations, and manifest as ecological pattern. It is therefore increasingly common to control for light in quantitative analyses (Kleypas et al. 1999). Similarly to our findings, Fricke & Meischner (1985) reported 1% PAR at 90-100 m depth in Bermuda; comparable to Okinawa (Yamazato 1972), and measurements from the northern Red Sea (Eyal et al. 2015). These publications, however, contain limited replication in space and time. We argue that in considering the role of light quantity and quality in structuring coral communities along large depth gradients, extensive measurements and replications are necessary to uncover the variability in water quality (e.g. resulting from desert flash-floods, algal blooms) (Kirk 2011).

### Light limitation and solar stress may control the shallow to mesophotic transition

Similarly to previous studies (Semmler et al. 2016, Laverick et al. 2017), our ecological analysis revealed two coral assemblages (clusters) between the surface and 100 m depth. The switch from shallow genera to mesophotic genera occurs around 50 m. Dekel is a particularly unusual site, with a shift in coral community structure between 30-40 m (Fig. 4). Cluster 1 comprised mainly ‘shallow’ genera. Depth ranges for cluster 2 were ‘deep’ but more varied than cluster 1 (Fig. 4S). Although our statistical analysis does not allow a clear statement with respect to coral assemblages differing as depth-specialists or generalist taxa, the taxa in each cluster do segregate loosely by depth (Fig. 4S).

The two assemblages exhibit different relationships with light (Fig. 4S). The quadrats most closely mirror the shallow assemblage above 0.29% surface UV. Below this, the shallow corals appear light limited and their DF values rapidly decline. The quadrats match the mesophotic assemblage most strongly at 7.5% surface PAR, with a decline above and below this number. This suggests that mesophotic taxa are only able to survive within a narrow band of light values. Below 7.5% PAR mesophotic taxa may be light limited, while above this value solar stress may limit the distribution of genera. The model displays a tail toward high light levels containing a few non-zero DF-values. As light readings were taken by casts in open water, they may not accurately reflect the light environment of a topographically complex reef. These non-zero DF values could have resulted from shaded photo-plots in shallow water. We are thus unable to disprove competing explanations for the loss of mesophotic taxa under higher light values. It may also be possible that competitive interactions with shallow taxa, which are better able to tolerate brighter conditions, restrict mesophotic taxa to darker reef patches. Both clusters are characterized by different morphological forms, which may reflect adaptations to physical conditions (Kahng et al. 2019).

It is worth noting that the asymptotic DF value for shallow assemblages in the quadrats is 0.3, twice the maximum value returned by the model for the mesophotic photo-plots. This can be explained by the average number of genera from each assemblage captured by the photo-plots. Plots assigned to cluster 1 on average captured 8.8 genera. Plots identified as cluster 2 captured 3.3 genera. This suggests that greater sampling effort is required at mesophotic depths in order to generate strong signals. These models indicate a physiological explanation for the community transitions found on coral reefs. In addition to solar radiation, temperature regimes too can influence corals (Cantin et al. 2010, Downs et al. 2013, Muir et al. 2015). In general, we found minimal changes in water temperatures across depth gradients and among sites throughout the year (Fig 5S).

### Seasonal algal blooms in the GoE/A modulate the light environment experienced by reefs, explaining the shallow-mesophotic reef transition

Tight relationships between physical and ecological patterns have been reported in previous studies (Labiosa et al. 2003, Boss and Behrenfeld 2010, Dishon et al. 2012). Similarly, we found that annual algal blooms in the GoE/A may limit the depth of shallow taxa. This ecological mechanism increases light attenuation, mainly in oligotrophic waters (Bricaud et al. 1998; Kirk 2011), and could also dictate the depths of mesophotic reefs (Figs. 5 and 6). The mesophotic coral *S. pistillata* was found to be a net producer of O_2_ only during the summer in the GoE/A (Nir et al. 2014). This coincides with the lowest Chl-*a* concentrations in the water column. At the height of the algal bloom, *S. pistillata* must rely on heterotrophy to satisfy its energy demand. Similar to other locations, the spring bloom in GoE/A occurs as a result of the mixed-layer depth (MLD) (Zarubin et al. 2017). Usually, the MLD maximum is reached in February-March, with substantial inter-annual variability in the maximum MLD between years (Zarubin et al. 2017). The cycles shown (Fig. 5) represent the relative difference in chlorophyll-*a* concentration expected in the water column over the course of a year. This seasonality was calculated from 14 years of monitoring data prior to the current study. Despite annual fluctuations, we can expect the benthic communities to respond to these long-term averaged signals. This can explain why March and June were selected by AIC as the best explanation of shallow and mesophotic community distributions, respectively, when considering PAR (Fig. 6). Modern tools, such as remote sensing, may provide information on water quality and light conditions in space and time. Phytoplankton, which play an important role in locally modulating the bleaching response (e.g. during episodes of heat stress), may be an influential factor in a mitigating way by reducing harmful light stress (Maina et al. 2008). Thus, it may not only influence bleaching response but also recruitment success at depth ranges. We encourage other researchers to consider the importance of seasonal events that potentially modify the light field as explanations for the patterns found in deep-reef communities elsewhere.

### Approaches for collecting water column irradiance data

We present multiple methods for acquiring light data to complement ecological surveys on MCEs. Spectral measurements with light meters (i.e. PRR-800, see methods) can provide PAR as well as UVR values in water and are common. Model-based remote-sensing approaches also exist, which can calculate surface PAR (Skirving et al. 2017). When combined with water quality data (Kd_(PAR)_), these models can give the photic depth of a site. In addition to Skirving’s method, NOAA has a global PAR product. This matches NOAA’s 0.05 degree (approx. 5 km) Geo-Polar Blended SST product. Information about the GOES Surface and Insolation Products (GSIP) available at: www.ospo.noaa.gov/Products/land/gsip/index_v3.html. However, despite these datasets being open source and providing important quantitative estimates of light qualities, measuring PAR alone is insufficient to also predict the attenuation of UVR (Dunne and Brown 1996).

When water-proof measuring equipment is unavailable, HydroLight software can be used to estimate light quality (available at: www.sequoiasci.com/product/hydrolight/). This platform computes in-water radiance as a function of depth, direction and wavelength (400-700nm in bandwidth 2nm). Other quantities of interest, such as the water-leaving radiance and remote-sensing reflectance, can also be obtained (Mobley 1999). HydroLight software could be combined with satellite data to produce meaningful light values, without the need for any measuring instrument (Mobley 1999, Lee et al. 2005).

If none of the above approaches are suitable, Secchi disks can be used to estimate the photic depth at a given location using Beer–Lambert’s law. Measurements need to be repeated over time to capture episodic events such as floods or algal blooms, and in space to capture variability between sites and depths. Such an approach is therefore more laborious than the satellite and modelling solutions.

### Why should we adopt an environmental definition of MCEs?

Our data have revealed variation in coral community structure in tandem with light conditions, across different sites (Fig. 6). Therefore, any definition of a mesophotic reef should have a flexible depth limit to account for the environmental conditions at a particular site. The correlation between the physiology and ecology data could be explained by light penetrating less deeply at specific sites (i.e. Dekel) (Figs. 4 & 6). Although there were no statistically significant differences in community structure across sites, there are coral depth distributions which are affected by changes in light. We should expect to find mesophotic species at shallow depths under certain light conditions (Kahng et al. 2010; Muir et al. 2015). i.e. *Alveopora* spp. flourish at 70 m at the clearest water sites (Katza and IUI), while no colonies were found at this depth at Dekel. In general, this genus, as well as *E. paradivisa* is distributed at deeper depths at sites with low Kd_(PAR)_ values (Fig. 6b). Thus, by providing a clearer definition to light-limited coral communities inhabiting MCEs, we may be able to better predict the implications of these effects on the recruitment, and post settlement survivorship of specific coral species.

### How variability in the light field affects coral species must be considered when projecting different futures for MCEs

MCEs are likely sensitive to fluctuations in the light field, induced by local disturbance (e.g. eutrophication, algae blooms). To support the persistence of mesophotic populations, we need to predict future population changes under different environmental scenarios. To do this accurately will require an understanding of the differing responses of coral groups (e.g. depth specialists or generalists, autotrophs or mixotrophs) to a variety of pressures, including the competitive influences between them (Laverick 2017). The ability to cope with changes in light quality, in combination with other stressors (e.g. thermal stress, acidification and pollution) is expected to vary among species (Bauman et al. 2013).

Photosynthetic taxa in the lower mesophotic may be the most affected by changes in light (Bongaerts et al. 2015) (Fig. 2. *E. paradivisa*, *Alveopora* spp., *L. fragilis).* This can be seen in the tight connection between different stony corals and the 1% PAR and UV limits (Fig. 2). The changing depth distributions for these species among sites appears connected to light (Fig. 6). These relationships may influence competition and the available settlement area on MCEs, under environmental change, and subsequently impair coral resilience to other stressors.

By combining ecological and physical (light) data it is possible to more accurately define a mesophotic and a shallow reef-community. This approach respects that an ecological pattern is dependent in part on physiological limitations. Further, it allows us to consider how reefs may respond to changing environmental conditions globally. Rapid changes in water quality may have a crucial effect, felt most keenly by depth-specialist taxa. These organisms may find themselves unable to keep pace with changes in the light regime, and so end up outside their preferred abiotic envelopes. Continuous ecological and physical monitoring is therefore needed to assess the health of sensitive mesophotic ecosystems. Consequently, when investigating the responses of corals on an ecological scale (e.g. changes in distribution and community structure), we suggest that experiments are needed to determine the environmental-physiological interactions that drive these patterns. By combining field and lab studies with light data, we can quantify the influence of light on the spatial distribution of stony corals.

## Acknowledgements

We would like to thank the Interuniversity Institute for Marine Sciences at Eilat (IUI) for making their facilities available to us. We are grateful to Motti Ohevia for construction of the underwater photography system and for technical help. This project was funded by the Israel Science Foundation (ISF) Grants No. 341/12 and 1191/16 to YL and by the European Union’s Horizon 2020 research and innovation program under the Marie Skłodowska-Curie post-doctoral grant agreement No. 796025 to GE.

## Data availability

Data available from:

NMP data:http://www.meteotech.co.il/EilatYam_data/ey_observatory_pier_download_data.asp

NOAA website: http://www.ospo.noaa.gov/Products/land/gsip/index_v3.html

Hydrolight website: http://www.sequoiasci.com/product/hydrolight/

---

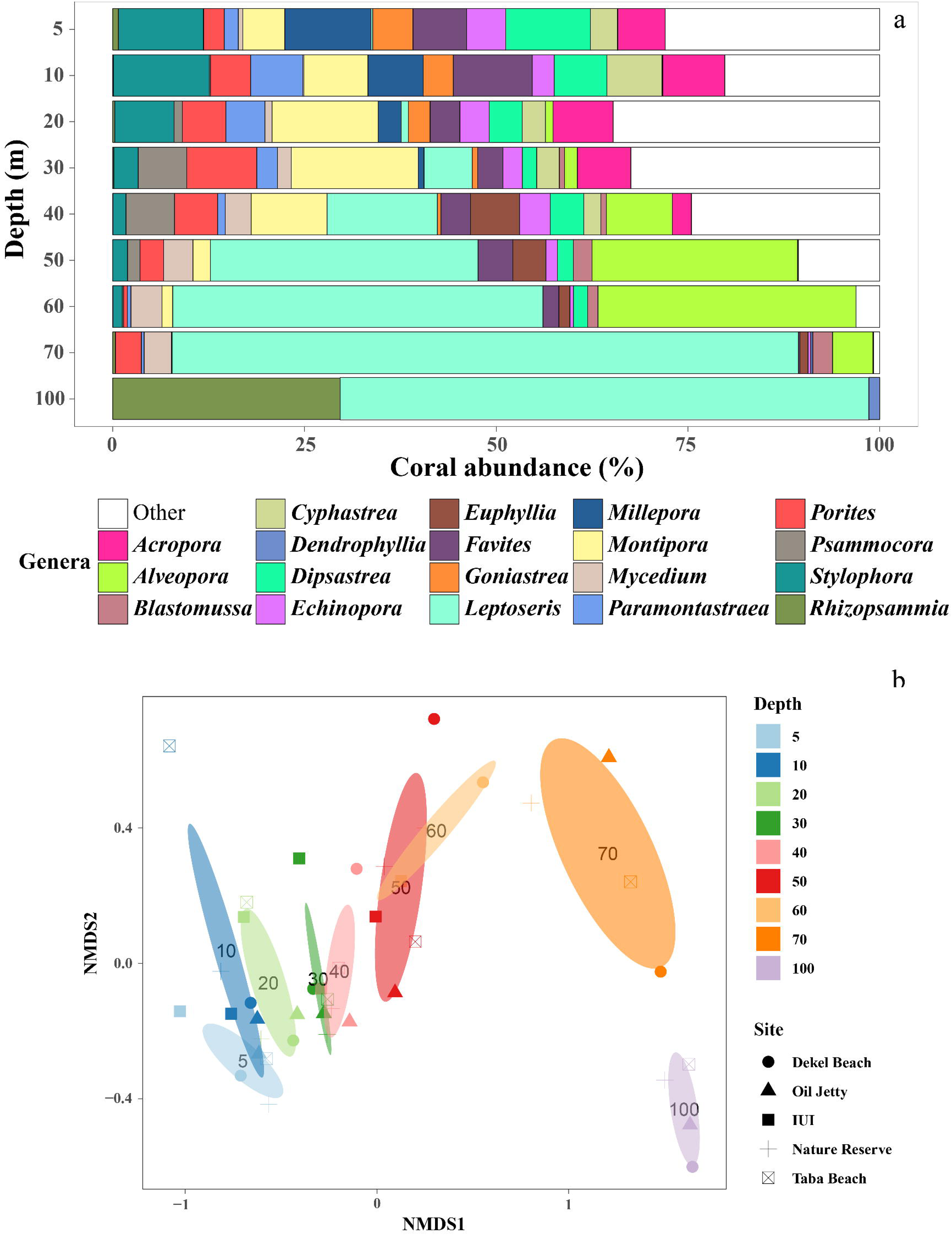

