## supplementary for "Light environment drives the shallow to mesophotic coral community transition"

*1. School of Zoology, George S. Wise Faculty of Life Sciences, Tel Aviv University, ISRAEL*

**Table 1S** Average dissimilarities for depths zone as determined by SIMPER and one-way ANOSIM pairwise tests. The overall model was significant (Global R = 0.802, p < 0.001, with 999 permutations). All the tests are significant. Shallow ≤ 50 m, Upper MCE = 50-70 m, Lower MCE ≥ 70 m.

| *Pairwise tests* | *Average dissimilarity* | *R statistic* | *p - value* |
|---|---|---|---|
| *Shallow - Upper MCE* | 65.01 | 0.72 | **0.001** |
| *Shallow - Lower MCE* | 94.18 | 1.00 | **0.001** |
| *Upper MCE - Lower MCE* | 74.03 | 0.76 | **0.001** |

**Table 2S** Pairwise comparisons of ANOSIM between the community compositions found at the 10 m depth interval transitions.

| *Pairwise groups* | *R statistic* | *p-value* |
|---|---|---|
| *5-10 m* | 0.076 | 0.183 |
| *10-20 m* | 0.124 | 0.143 |
| *20-30 m* | 0.196 | 0.159 |
| *30-40 m* | -0.004 | 0.484 |
| *40-50 m* | 0.404 | **0.024** |
| *50-60 m* | -0.005 | 0.464 |
| *60-70 m* | 0.593 | 0.057 |
| *70-100 m* | 0.74 | **0.029** |
| *Global test* | 0.76 | **0.001** |

**Table 3S.** AIC values used in model selection. The month of light data used when comparing models explaining DF values with either depth, PAR, or UV, is underlined and in bold.

| *Month* | *AIC: PAR* | | *AIC: UV* | |
|---|---|---|---|---|
| | ***Cluster 1*** | ***Cluster 2*** | ***Custer 1*** | ***Cluster 2*** |
| *January* | *NA* | *NA* | *NA* | *NA* |
| *February* | -994 | -2886 | -1080 | -2790 |
| *March* | **-1034** | -2821 | *NA* | *NA* |
| *April* | -977 | -2960 | -1085 | -2791 |
| *May* | -1005 | -3093 | **-1099** | -2804 |
| *June* | -983 | **-3249** | -1085 | -2824 |
| *July* | *NA* | *NA* | *NA* | *NA* |
| *August* | -987 | -3191 | -1086 | **-2845** |
| *September* | -629 | -2106 | -1081 | -2805 |
| *October* | *NA* | *NA* | -1080 | -2790 |
| *November* | -995 | -2882 | -1086 | -2776 |
| *December* | -994 | -2905 | -1089 | -2789 |

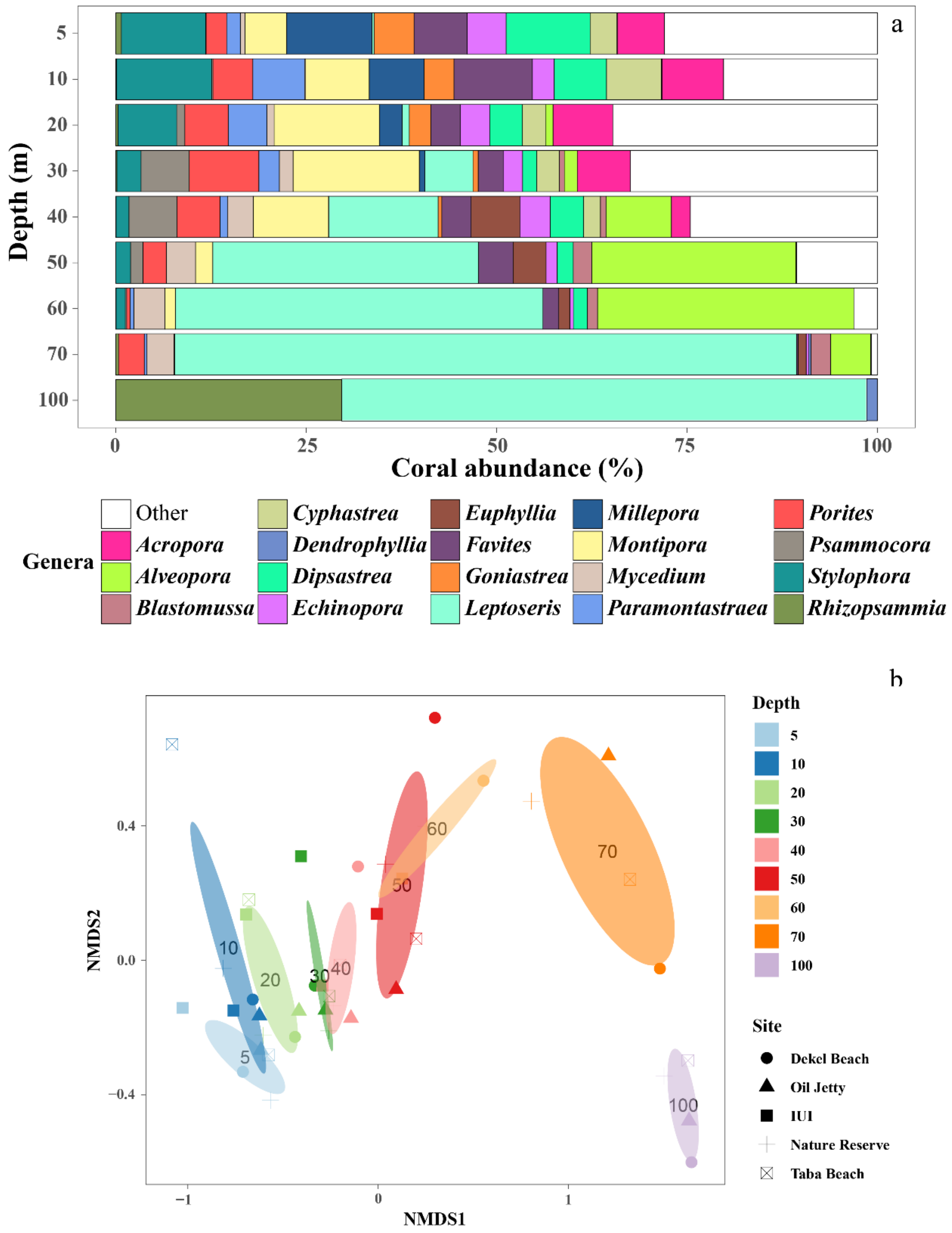


**Fig. 1S.** (a) Total cumulative percentage of the relative abundance of the 19 most common scleractinian coral genera along a depth gradient at the five survey sites. (b) NMDS ordination using scleractinian coral community composition from a depth gradient ranging from 5 to 100 m depth. Each colour represents a particular depth, and each dot the community composition at this depth for a given site, based on Bray-Curtis dissimilarity matrices (2D, stress = 0.07). Polygons are SE of depth groups.

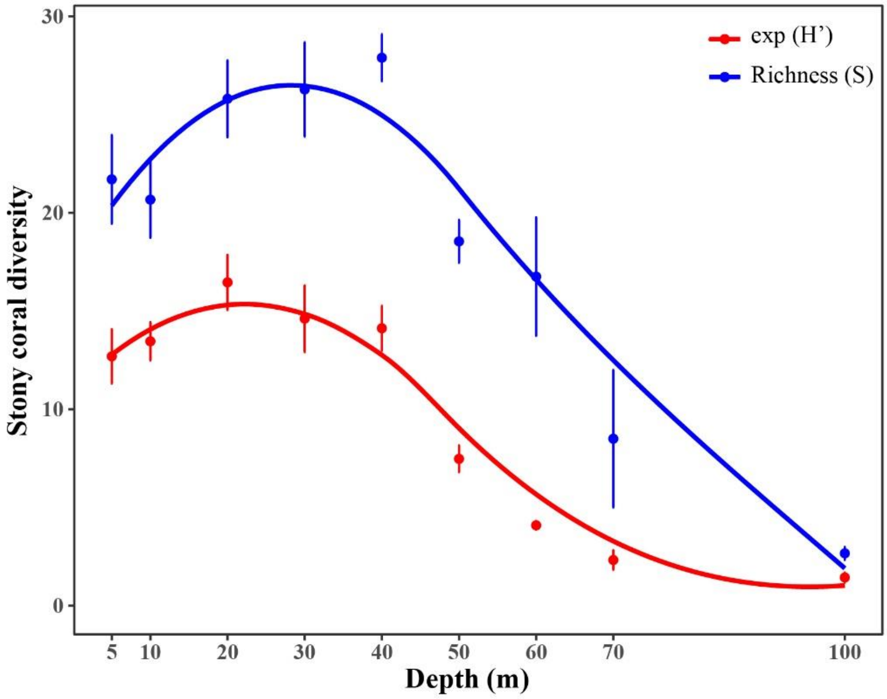


**Fig. 2S.** Stony corals Shannon index Hill number (exp (H')) (red) and Species Richness (S) (blue), along surveys depth gradient. Error bars represents ±SE.

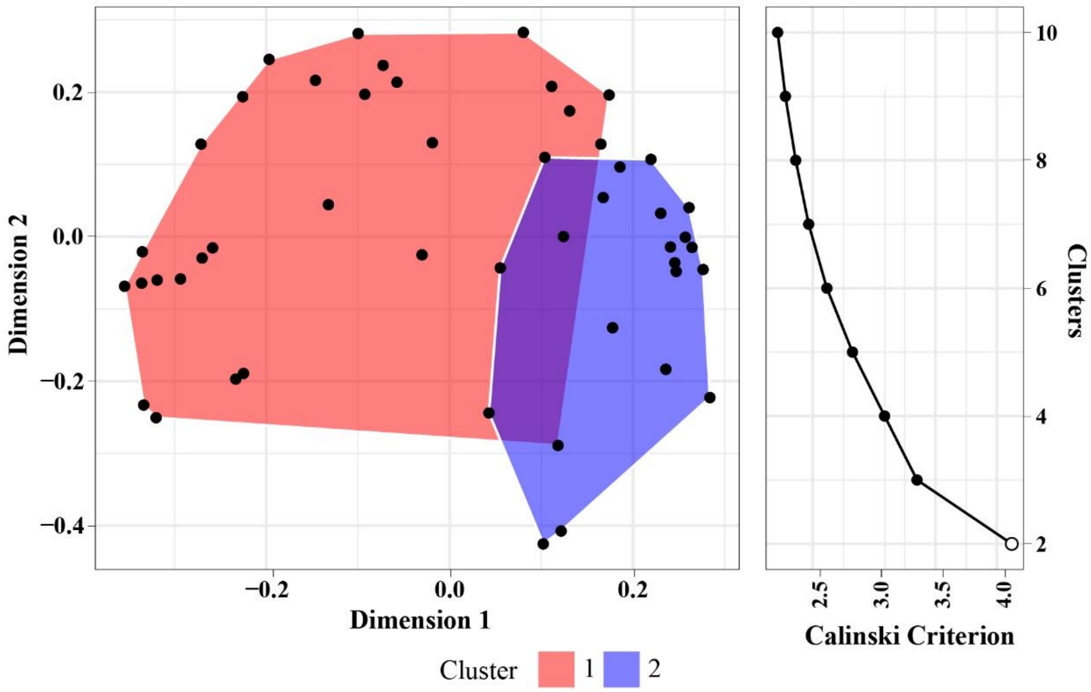


**Fig. 3S.** Principal Co-ordinate analysis of quadrat data was used to identify assemblages of co-occurring Scleractinia genera with K means clustering (left). Polygons show the degree of overlap between clusters in taxon space. The Calinski criterion of proposed number of clusters to fit to the data is shown to the right. The optimal number of clusters was considered to be the number that maximised the Calinski criterion, marked as a hollow point.

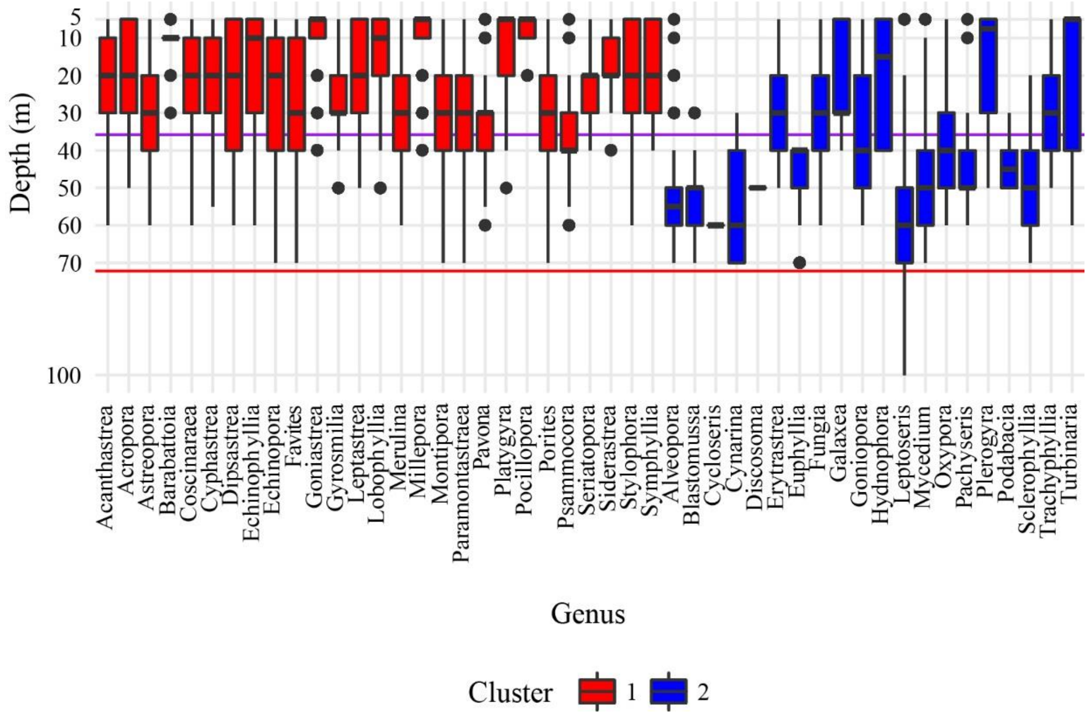


**Fig. 4S.** Boxplots show the depth ranges of 47 genera generated from pooled data of all sites. Boxes are coloured according to assemblage identified during PCoA analysis. Cluster 1 (red) and genera from cluster 2 (blue). 'Whiskers' of the boxplot extend 1.5x the interquartile range, or to the most extreme observation, whichever is smaller. Points are outliers beyond 1.5x the interquartile range from the median. The density curve on the right of the plot shows the distribution of all observed coral colonies with increasing depth. Horizontal black lines are the mean 1% UV (purple) and mean 1% PAR (red) limits across all sites.

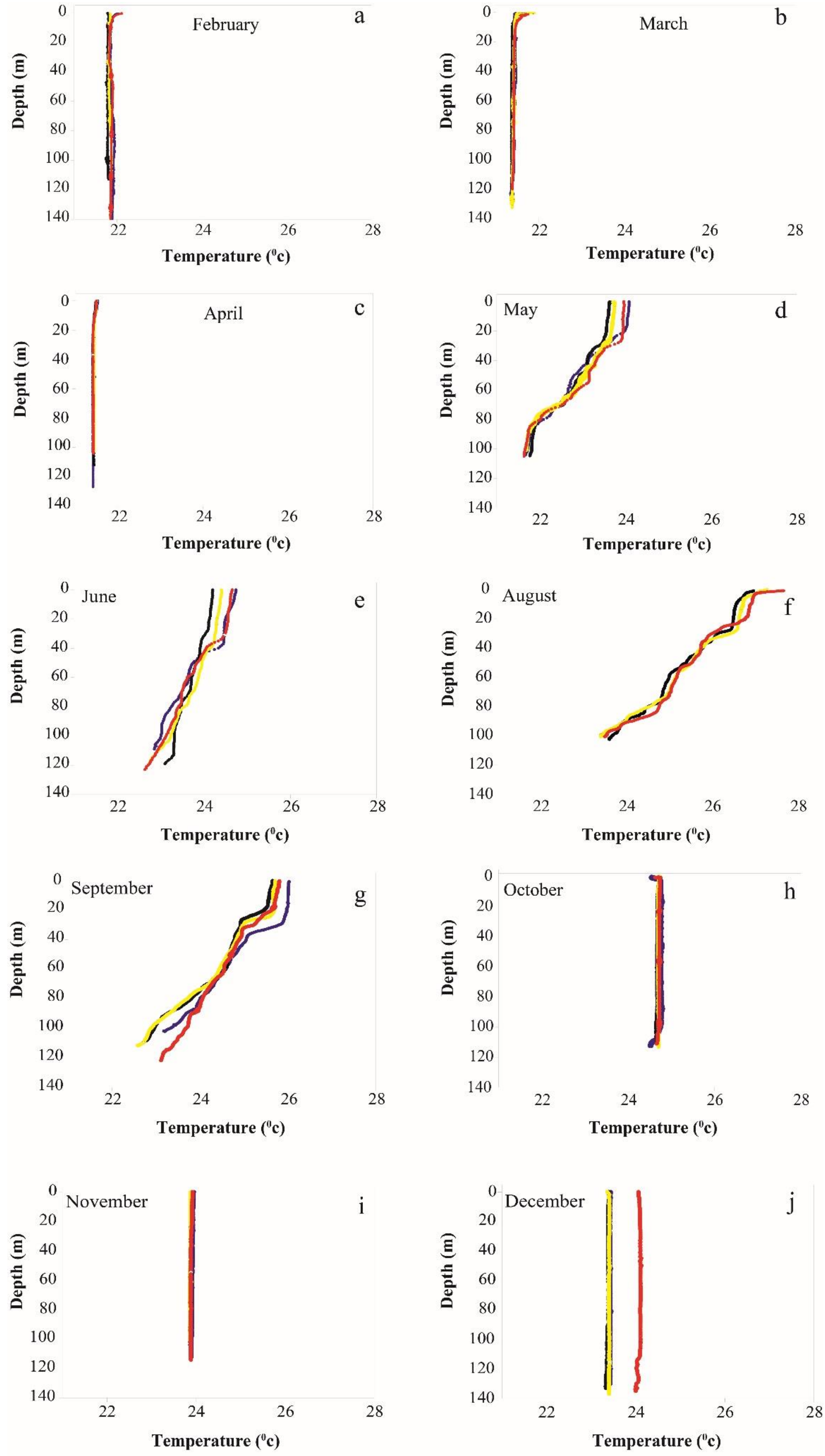


**Fig. 5S.** Water column temperature at the four sites – Dekel Beach (black), Katza (yellow), IUI (red) and Taba (blue), from February to December between 2014-2015.
